# Disturbance-Diversity Relationships of Microbial Communities Change Based on Growth Substrate

**DOI:** 10.1101/2023.08.25.554838

**Authors:** Don Q. Hoang, Lindsay R. Wilson, Andrew J. Scheftgen, Garret Suen, Cameron R. Currie

## Abstract

Disturbance events can impact ecological community dynamics. Understanding how communities respond to disturbances, and how those responses can vary, is a challenge in microbial ecology. In this study, we grew a previously enriched specialized microbial community on either cellulose or glucose as a sole carbon source, and subjected them to one of five different disturbance regimes of varying frequencies ranging from low to high. Using 16S rRNA gene amplicon sequencing, we show that community structure is largely driven by substrate, but disturbance frequency affects community composition and successional dynamics. When grown on cellulose, bacteria in the genera *Cellvibrio*, *Lacunisphaera*, and *Asticaccacaulis* are the most abundant microbes. However, *Lacunisphaera* is only abundant in the lower disturbance frequency treatments, while *Asticaccaulis* is more abundant in the highest disturbance frequency treatment. When grown on glucose, the most abundant microbes are two *Pseudomonas* sequence variants, and a *Cohnella* sequence variant that is only abundant in the highest disturbance frequency treatment. Communities grown on cellulose exhibited a greater range of diversity (0.67-1.99 Shannon diversity and 1.38-5.25 Inverse Simpson diversity) that peak at the intermediate disturbance frequency treatment, or 1 disturbance every 3 days. Communities grown on glucose, however, ranged from 0.49-1.43 Shannon diversity and 1.37-3.52 Inverse Simpson with peak diversity at the greatest disturbance frequency treatment. These results demonstrate that the dynamics of a microbial community can vary depending on substrate and the disturbance frequency, and may potentially explain the variety of diversity-disturbance relationships observed in microbial ecosystems.

**Abstract Importance:** A generalizable diversity-disturbance relationship (DDR) of microbial communities remains a contentious topic. Various microbial systems have different DDRs. Rather than finding support or refuting specific DDRs, we investigated the underlying factors that lead to different DDRs. In this study, we measured a cellulose-enriched microbial community’s response to a range of disturbance frequencies from high to low, across two different substrates: cellulose and glucose. We demonstrate that the community displays a unimodal DDR when grown on cellulose, and a monotonically increasing DDR when grown on glucose. Our findings suggest that the same community can display different DDRs. These results suggest that the range of DDRs we observe across different microbial systems may be due to the nutritional resources microbial communities can access and the interactions between bacteria and their environment.

## Introduction

Disturbance ecology investigates foundational questions of how systems and organisms respond to changing environments. Traditionally, disturbances are defined as discrete events that remove biomass directly or indirectly through displacement or mortality ^1,2^. Fires, floods, and volcanic eruptions are classic examples of disturbances that change community composition by directly impacting species or altering the environment ^3,4^. Early theoretical consideration of disturbance on community ecology include the Intermediate Disturbance Hypothesis (IDH), which predicts that the diversity-disturbance relationship (DDR) follows a “hump-backed”, or unimodal curve ^5^. Support for the IDH has been mixed. Experimental measurements of the DDR for different systems has revealed a variety of trends, including both positive and negative monotonic, unimodal, bimodal, and several nonsignificant DDRs ^6^. Recent frameworks of disturbance theory accommodate vastly different spatiotemporal scales between systems, and disentangle disturbance events and impacts ^7,8^.

The advent of high-throughput sequencing has widened the scope of questions that microbial ecology can ask, including research that investigates how disturbance impacts microbial communities. Researchers have studied disturbances in several different systems including marine sediment ^9^, soil bacterial ^10^ and soil fungal communities ^11^, and wastewater communities^12^. Microbial systems also display a variety of DDRs, which suggests that rather than trying to support or reject specific DDRs, researchers can better understand disturbance ecology by investigating the underlying factors that lead to different DDRs. Given the vast differences in systems between these studies, it is difficult to determine what specific factors lead to differing responses to a disturbance. Although we know that microbial community responses to disturbances can vary, whether the same community can exhibit different responses to the same disturbance and what factors would cause those differences, is relatively underexplored. Moreover, understanding what factors influence responses to a disturbance event is important for predictive power in studying microbial communities.

To address this gap in knowledge, we examined the effects of disturbance on a bacterial community enriched from the refuse pile of the leaf-cutter ant *Atta colombica* that had previously been passaged in the lab on minimal media and cellulose by Lewin et al. ^13,14^. Leaf-cutter ant refuse piles are composed of discarded plant biomass that has been partially degraded by the ants’ mutualistic fungal cultivar, *Leucoagaricus gongylophorus* ^15^. Previous work has demonstrated that these refuse piles are enriched with plant-biomass degrading microbes ^16,17^. Focusing on bacterial communities derived from leaf-cutter ant refuse piles, Lewin et al. experimentally evolved cellulose-degrading bacterial communities and investigated their compositional dynamics and cellulolytic abilities ^13,14^. During each passage, a portion of the community was aliquoted into a new test tube containing fresh minimal media and a new strip of cellulose. These serial transfer events are analogous to disturbance events, as it is a species-independent method of biomass reduction and provides the “survivors” with a replenished ecosystem. This method of proxying disturbance through removing cells has been used in other studies ^18^.

Lewin’s community was enriched on cellulose, a recalcitrant crystal of β-1,4-linked glucose molecules. Cellulose is insoluble in water, and must be cleaned into cellobiose or glucose in order to be transported into a cell. Cellulase genes have limited distribution in bacteria, but β-glucosidases, which cleave cellobiose into glucose, are more widespread ^19^. Thus, cellulolytic and noncellulolytic microbes compete for cellobiose – and these interactions may impact the community’s composition ^20,21^. Lewin et al. 2022 evaluated successional dynamics in this microcosm by measuring the relative abundance of 16S rRNA genes every day for a week and found that a *Cellvibrio* operational taxonomic unit (OTU) was more abundant up to 48-72 hours, before other OTUs became more abundant ^14^. This finding suggests that a cellulose degrader must proliferate and produce cellulases before noncellulolytic opportunists can take advantage of liberated cellobiose or metabolic byproducts.

Our goal was to understand how substrate complexity interacts with disturbance frequency to shape community diversity. We hypothesize that diversity maximizes on a simple substrate (glucose) at higher disturbance frequencies but maximizes on a complex substrate (cellulose) at lower disturbance frequencies. We reasoned that on a simple substrate with low disturbance, competition exclusion would be a stronger driving force for community assembly while more frequent disturbances would disrupt competitive microbes from establishing. Conversely, on complex substrates, the ability to use the substrate would be a more important driving force. Since cellulases are phylogenetically limited in distribution ^19^, we hypothesize that the bacteria that initially grow will be those that can degrade cellulose similar to what Lewin et al. 2022 observed ^14^. Frequent disturbances should select for bacteria that can directly use cellulose. As cellulose is degraded into cellobiose, those molecules enrich the surrounding media and feed non-degraders. Thus, at infrequent disturbances non-degraders can grow making the community taxonomically richer.

To test our hypotheses that diversity maximizes on glucose at high disturbance frequencies and maximizes on cellulose at lower disturbance frequencies, we subjected Lewin et al.’s cellulose-enriched community to two substrate treatments: minimal media supplemented with either glucose or cellulose. Each substrate was then subjected to five disturbance frequencies: passage every 1, 2, 3, 5, or 7 days. At the end of their assigned disturbance regime, we expanded the communities into multiple tubes of their respective substrate and destructively sampled over the course of one week. We then extracted DNA from these samples for 16S rRNA gene amplicon Illumina-based sequencing. Next, we analyzed these sequences to determine community composition and measured diversity. By comparing the same disturbance frequency between substrate complexities, we can evaluate how community diversity is affected by the interaction between disturbances and resources.

## Methods

### Enrichment on cellulose

The bacterial community used here was previously enriched on cellulose as reported by Lewin et al ^13,14^. Briefly, approximately 3 mg of refuse dump originating from *A. colombica* refuse piles was added to test tubes containing 5 mL of M63 minimal media, and a 1×10 cm strip of Whatman Grade 1 cellulose filter paper (GE Healthcare Life Sciences, Pittsburgh, PA). As the filter paper was the only carbon source, it selected for a cellulolytic community. The tubes were grown aerobically with shaking at 30 °C. Once the filter paper broke, the test tube was vortexed, and 200 μL of the culture was transferred into a new test tube containing fresh M63 media and a new strip of filter paper. This transferring process was repeated each time the filter paper broke. These communities were previously reported at 20 transfers ^13^, and 60 transfers ^14^. For this work, we used community “3A” from Lewin et al. 2022.

### Media Used

M63 minimal media was used for all experiments. This media was modified with different carbon sources, depending upon the treatment. For glucose treatments, 12.5 mL of 40% filter sterilized glucose was added to each liter of M63 media. For cellulose treatments, 5 grams of cellulose powder per 1 liter of M63 was added. For tubes that used cellulose paper, 1 strip of 1 x 10 cm Whatman Grade 1 cellulose filter paper (GE Healthcare Life Sciences, Pittsburgh, PA) was added to a test tube containing 5mL of M63 media. Media recipes can be found in Supplementary Table 1.

### Disturbance experiment

To investigate how substrate complexity and disturbance frequency interact to shape community dynamics, we grew Lewin’s cellulose enriched community from a freezer stock of transfer #73 in M63 media supplemented with cellulose filter paper with shaking at 30 °C. Once the community broke the cellulose filter paper, it was scaled up into a 250 mL culture of M63 media and cellulose filter paper. After four days of growth (when we observed visible cellulose paper degradation) the community was split into test tubes representing 10 different treatments (Figure 1). Communities were grown in 5 mL of M63 minimal media supplemented with either glucose or cellulose. These two media treatments were further divided into five disturbance treatments: every 1, 2, 3, 5, or 7 days test tubes were vortexed to homogenize the community and 200 μL of the culture was used to inoculate a new tube of fresh media. Each treatment had five technical replicates. Each disturbance was carried out 10 times. That is, tubes in the 1 day disturbance treatment were transferred every day for 10 days and tubes in the 7 day disturbance treatment were transferred every week for 10 weeks. We refer to these disturbance treatments as 1/n days. For example, treatments that were passaged every 3 days will be referred to as “1/3 days,” indicating 1 disturbance per 3 days. At the end of their respective disturbance regime, each community was used to inoculate 7-10 tubes which were destructively sampled every day for one week to evaluate any compositional dynamics. Those tubes were then destructively sampled each day for one week. At sampling, the bacterial culture was spun down and the pellet and supernatant were separated. Cell pellets were resuspended in Zymo DNA/RNA shield (Zymo Research, Irvine, CA, USA) and stored at −80°C until DNA extraction.

**Figure 1.**
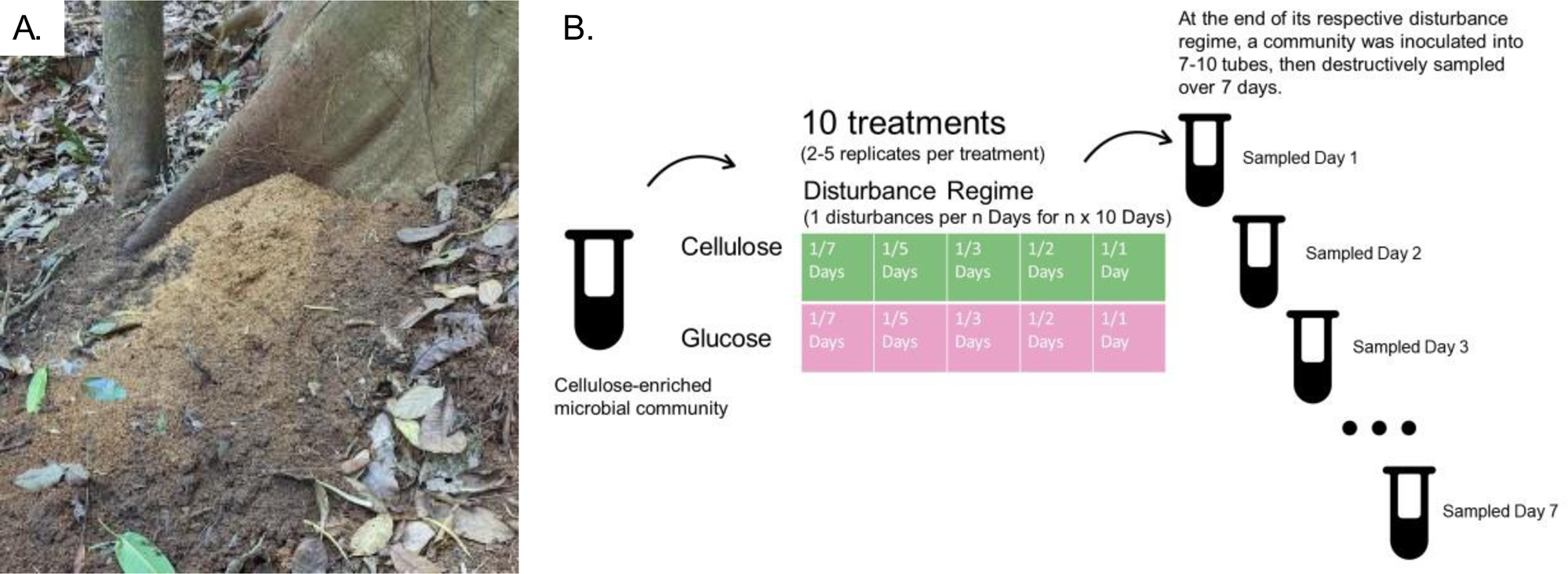
A. Image of A. colombica refuse dump. These piles are predominantly composed of partially degraded plant biomass, removed from the bottom of fungus gardens by worker ants. B. Experimental setup. The cellulose-enriched community was used as the starter inoculum for the growth of microbial communities exposed to ten treatments. Samples were grown in either cellulose or glucose M63 minimal media, then subjected to one of five disturbance regimes. At the end of their respective disturbance regime, communities were used to inoculated 7-10 tubes (containing their respective substrate) and those tubes were destructively sampled every day for seven days.

### Extraction and Sequencing

We extracted DNA for 16S rRNA gene amplicon sequencing. Samples were extracted using QIAGEN DNeasy PowerSoil kits (QIAGEN, Hilden, Germany) following the manufacturer’s instructions.

For 16S rRNA gene-based community profiling, the V3 to V4 regions of the 16S rRNA gene were amplified using bacteria-specific primers ^22^. Each reaction contained 50 ng DNA, 0.4 μM forward primer, 0.4 μM reverse primer, 12.5 μL 2X HotStart ReadyMix (KAPA Biosystems, Wilmington, MA, USA), and water to a final volume of 25 μL. Polymerase chain reaction (PCR) was performed using a Bio-Rad S1000 thermocycler (Bio-Rad Laboratories, Hercules, CA, USA). Cycling conditions began with initial denaturation at 95 °C for 3 minutes, followed by 25 cycles of 95 °C for 30 seconds, 55 °C for 30 seconds, and 72 °C for 30 seconds, and a final extension at 72 °C for 5 minutes. We included controls using sterile water in place of DNA to ensure there was no contamination during PCR. PCR products were purified using gel extraction from a 1.0% low-melt agarose gel (National Diagnostics, Atlanta, GA, USA) with a ZR-96 Zymoclean DNA Recovery Kit (Zymo Research, Irvine, CA, USA) and DNA was quantified using a Qubit Fluorometer and Qubit Kit (Invitrogen, Carlsbad, CA, United States). Samples were equimolarly pooled with 10% PhiX control DNA and sequenced on an Illumina MiSeq using a MiSeq 2 × 250 v2 kit (Illumina, Inc.). Due to the number of samples, sequencing was performed across two sequencing runs.

Reads were processed using DADA2 ^23^ in R version 4.2.1 ^24^ with taxonomy assignment using the SILVA V138.1 references database ^25,26^. Sequencing yielded 11,670,039 sequences, and denoising, merging, and removal of chimeric sequences resulted in 10,499,700 sequences. Samples had an average of 29,410.92 ± 33,930.7 sequences, and an average of 18.41 ± 12.62 ASVs. Of the 357 samples we sequenced, 302 were used for analysis. We removed negative control samples, samples with low reads (less than 1000 reads), and five samples that we suspect were mislabeled (Supplemental Fig. 1). Analysis was performed using the phyloseq ^27,28^ and vegan packages ^29,30^ in R. The Shannon Index, Inverse Simpson Index, Pielou’s Index, and Menhinick’s Index were used to assess alpha-diversity. A Bray-Curtis dissimilarity matrix was calculated to assess beta-diversity. Scripts for read processing, diversity analysis, and figure generation can be found at (https://github.com/donnyhoang/cellulose_disturbance).

### Data availability

Amplicon sequencing data have been uploaded to the NCBI databases under BioProject number PRJNA1008240. Supplemental File 1 contains individual accession numbers and relevant metadata for each sample.

## Results

### Community composition varies between treatments

Communities grown on different substrates were found to be dominated by different microbes (Fig. 2). Across those treatments with cellulose as the only carbon source samples, the most abundant microbes include *Cellvibrio*, *Lacunisphaera*, and *Asticaccaulis*, while two *Pseudomonas* amplicon sequence variants (ASVs) dominated samples with glucose as the only carbon source. The two *Pseudomonas* ASVs that dominate our glucose samples are not abundant across our cellulose samples, with a mean of 1.15% ± 1.77% and 4.72% ± 6.99% relative abundance. Similarly, the three most abundant ASVs in our cellulose samples were not abundant across the glucose samples. *Cellvibrio* (0.12% ± 0.34%), *Lacunisphaera* (0.019% ± 0.048%), and *Asticaccaulis* (0.15% ± 0.31%) are all under 1% relative abundance of the glucose samples.

**Figure 2.**
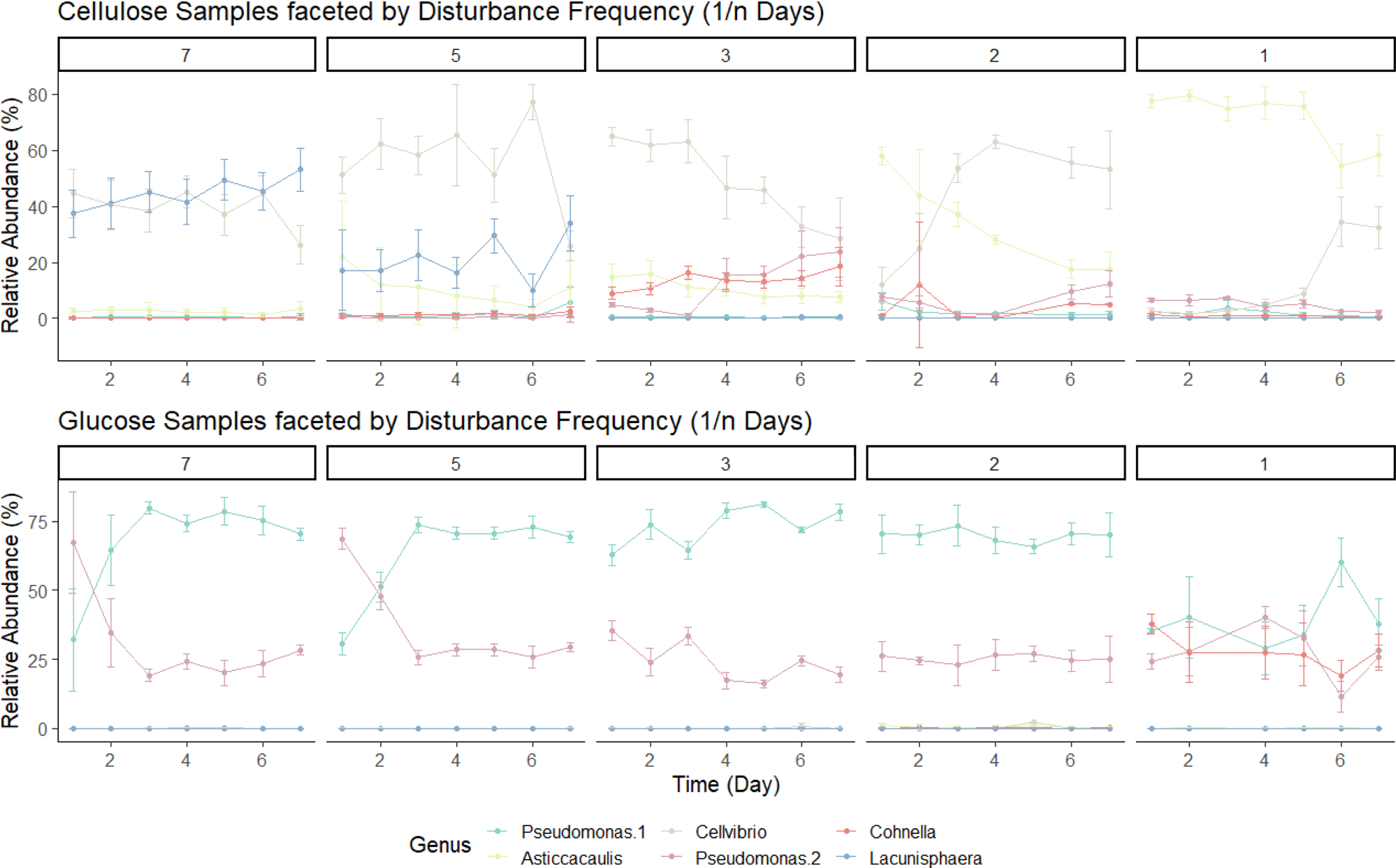
Line graph of community composition based on relative abundance of the 16S rRNA gene amplicon sequence of the 6 most abundant ASVs and their assigned genus. Plots are facetted by disturbance frequency (1/n days).

Disturbance frequency also impacted community composition when communities are grown on cellulose. In the cellulose substrate treatments, communities were dominated by *Asticaccaulis* (71.78% ± 10.66%) in the highest disturbance frequency (1/1 days). *Lacunisphaera* and *Cellvibrio* were present at lower relative abundance (0.02% ± 0.03 and 11.75% ± 13.94% respectively) in highest disturbance frequency treatment, 1 per 1 day. At the lowest disturbance frequency (1/7 days) the community was dominated by *Lacunisphaera* (44.79% ± 8.65%) and *Cellvibrio* (39.31% ± 9.2%) while *Asticaccaulis* was much lower (2.52% ± 1.64%).

Disturbance frequency does not impact the community composition of our glucose samples as strongly as it did in the cellulose treatment. Two Pseudomonas ASVs dominate nearly all the glucose samples. One *Pseudomonas* ASV ranged from 39.37% to 73.1% relative abundance, and the second *Pseudomonas* ranges from 24.13% to 35.4% relative abundance. A third ASV belonging to the genus, *Cohnella*, becomes abundant in the highest disturbance frequency treatment at 26.94% ± 9.2% relative abundance.

Disturbance frequency also impacts community dynamics. When cellulose samples were grown in more frequent disturbance treatments, 1/3 days 1/2 days and 1/1 days, there was a clear change over time in community composition. In the 1/3 frequency treatment, *Cellvibrio* was initially abundant (64.9% ± 3.32% on Day 1 to 28.3% ± 14.59% on Day 7), but gives way to *Pseudomonas* and *Cohnella* (23.59% ± 8.97% and 18.46% ± 6.84% respectively on Day 7). In the 1/2 days frequency treatment, *Asticcacaulis* is initially abundant (58% ± 3.23% on Day 1 to 17.33% ± 6.23% on Day 7) before *Cellvibrio* increases in abundance as the week progresses (12.01% ± 6.23% on Day 1 to 53.20% ±13.95% on Day 7). The 1/1 days frequency treatment showed a similar trend to the 1/2 days frequency treatment, where *Asticcacaulis* was dominant across the week of sampling (77.57% ±2.32% on Day 1 to 58.19% ± 7.27% on Day 7) and *Cellvibrio* starts low but increases in abundance (2.63% ± 0.95% on Day 1 to 32.45% ± 7.47% on Day 7). The cellulose substrate samples subjected to lower disturbance frequency treatments (1/7 days and 1/5 days) did not have a clear successional pattern.

Glucose samples largely did not show change in community composition across the week we sampled, with most of the community dominated by two *Pseudomonas* ASVs. However, the disturbance frequency treatments 1/7 days and 1/5 days showed some dynamics. In the 1/7 days disturbance frequency treatment, one *Pseudomonas* ASV increases (32.08% ± 18.53 on Day 1 to 70.28% ± 2.1% on Day 7) while a second *Pseudomonas* ASV decreases (67.23% ± 18.37% on Day 1 to 28.3% ± 1.75% on Day 7) in relative abundance. The 1/5 days treatment showed a similar trend where one *Pseudomonas* ASV increases (32.08% ± 18.53% on Day 1 to 69.28% ± 2.04% on Day 7) and the second *Pseudomonas* ASV decreases (67.23% ± 18.37% on Day 1 to 29.43% ± 1.69% on Day 7).

### Communities cluster by growth substrate

To better quantify the differences between our samples, we calculated Bray-Curtis dissimilarities across samples and visualized this as an NMDS plot (Fig 3., Supplemental Fig. 4). To test whether there is a significant difference between sample groups, we used the anosim() function in the R package vegan. We found that samples were most dissimilar based on Substrate (ANOSIM R = 0.96, p-value = 0.001). Samples separate along the NMDS1 axis based on substrate, and form two distinct clusters. Additionally, samples grown on cellulose display greater within treatment variation compared to glucose samples, as represented by a wider range across each NMDS axis. Within the two main clusters, samples also group by their disturbance frequency treatment. For example, samples grown in cellulose and subjected to disturbance frequency of 1/1 day group together, and samples grown in glucose and subjected to disturbance frequency of 1/1 day group together. However, 1/1 day disturbance frequency treatment samples do not group together nor align along either NMDS axis.

**Figure 3.**
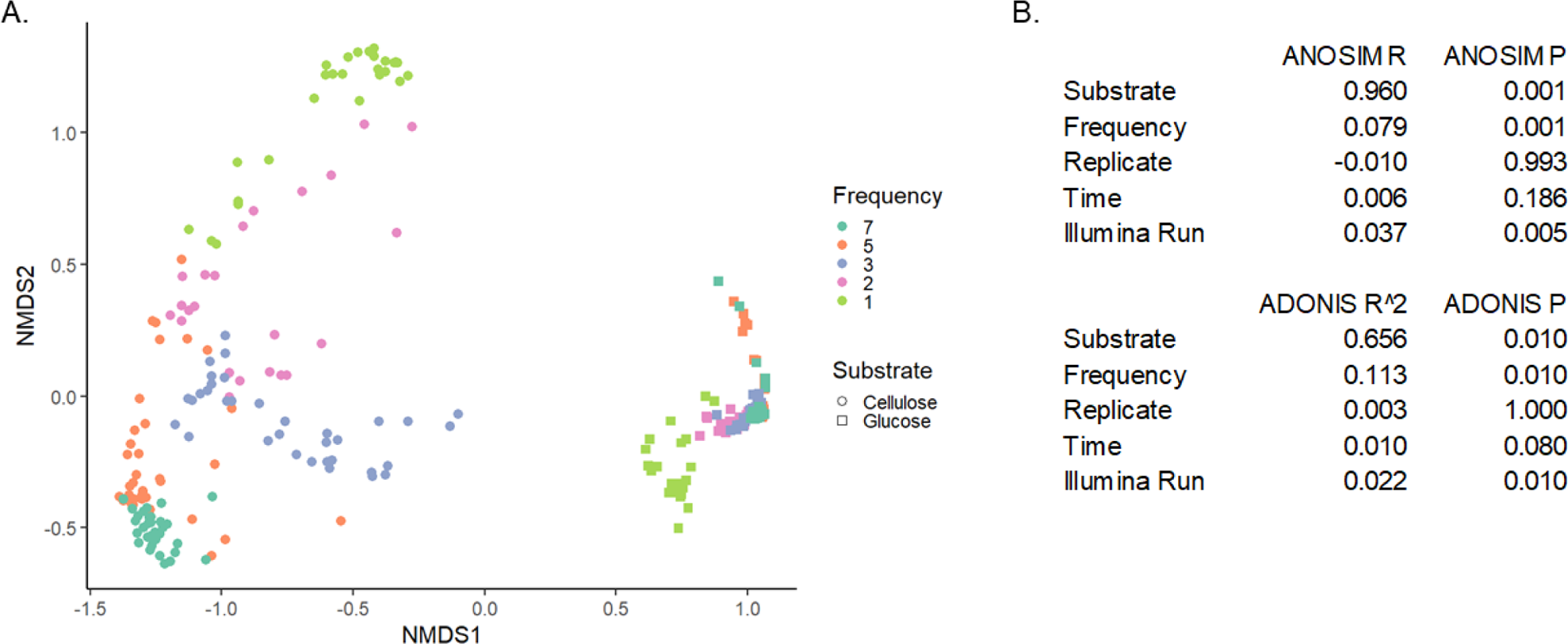
A. NMDS plot of community composition. Dissimilarity matrix was calculated using Bray-Curtis dissimilarity. Communities grown in cellulose are shown as circles, and communities grown in glucose are shown as triangles. Disturbance frequency is marked by color. B. Table of ANOSIM and ADONIS tests of recorded factors that may contribute to variance.

**Table 1.** Table of pairwise comparisons between Cellulose 1/3 treatment to other Cellulose disturbance frequency treatments, and Glucose 1/1 treatment to other Glucose disturbance frequency treatments.

| Diversity Index | Reference | Comparison | p-value | p.signif |
| --- | --- | --- | --- | --- |
| Shannon | Cellulose 1/3 | Cellulose 1/7 | 5.46E-06 | **** |
| Shannon | Cellulose 1/3 | Cellulose 1/5 | 4.93E-05 | **** |
| Shannon | Cellulose 1/3 | Cellulose 1/2 | 0.004326 | ** |
| Shannon | Cellulose 1/3 | Cellulose 1/1 | 6.35E-15 | **** |
| InverseSimpson | Cellulose 1/3 | Cellulose 1/7 | 0.0017 | ** |
| InverseSimpson | Cellulose 1/3 | Cellulose 1/5 | 0.00689 | ** |
| InverseSimpson | Cellulose 1/3 | Cellulose 1/2 | 0.003481 | ** |
| InverseSimpson | Cellulose 1/3 | Cellulose 1/1 | 4.18E-10 | **** |
| Shannon | Glucose 1/1 | Glucose 1/7 | 1.88E-21 | **** |
| Shannon | Glucose 1/1 | Glucose 1/5 | 8.33E-19 | **** |
| Shannon | Glucose 1/1 | Glucose 1/3 | 6.74E-20 | **** |
| Shannon | Glucose 1/1 | Glucose 1/2 | 8.73E-17 | **** |
| InverseSimpson | Glucose 1/1 | Glucose 1/7 | 5.03E-14 | **** |
| InverseSimpson | Glucose 1/1 | Glucose 1/5 | 1.13E-12 | **** |
| InverseSimpson | Glucose 1/1 | Glucose 1/3 | 6.62E-14 | **** |
| InverseSimpson | Glucose 1/1 | Glucose 1/2 | 9.12E-13 | **** |

### Diversity-disturbance relationships differ based on substrate

To evaluate how substrate complexity and disturbance frequency interact to affect community diversity, we measured the Shannon diversity and Inverse-Simpson diversity of our samples. We found that diversity changes across disturbance frequency, but this pattern differs depending on the substrate the community was grown in. Samples grown in cellulose increased in diversity (Shannon and Inverse Simpson indices), with a Shannon’s index of 1.99 for 1/3 disturbance frequency, but decreased as disturbances became more frequent (Fig. 4). Samples grown in glucose had the lowest measured diversity at low disturbance frequency, but diversity increased with disturbance frequency, achieving a Shannon’s index of 1.43 at 1/7 disturbance frequency.

**Figure 4.**
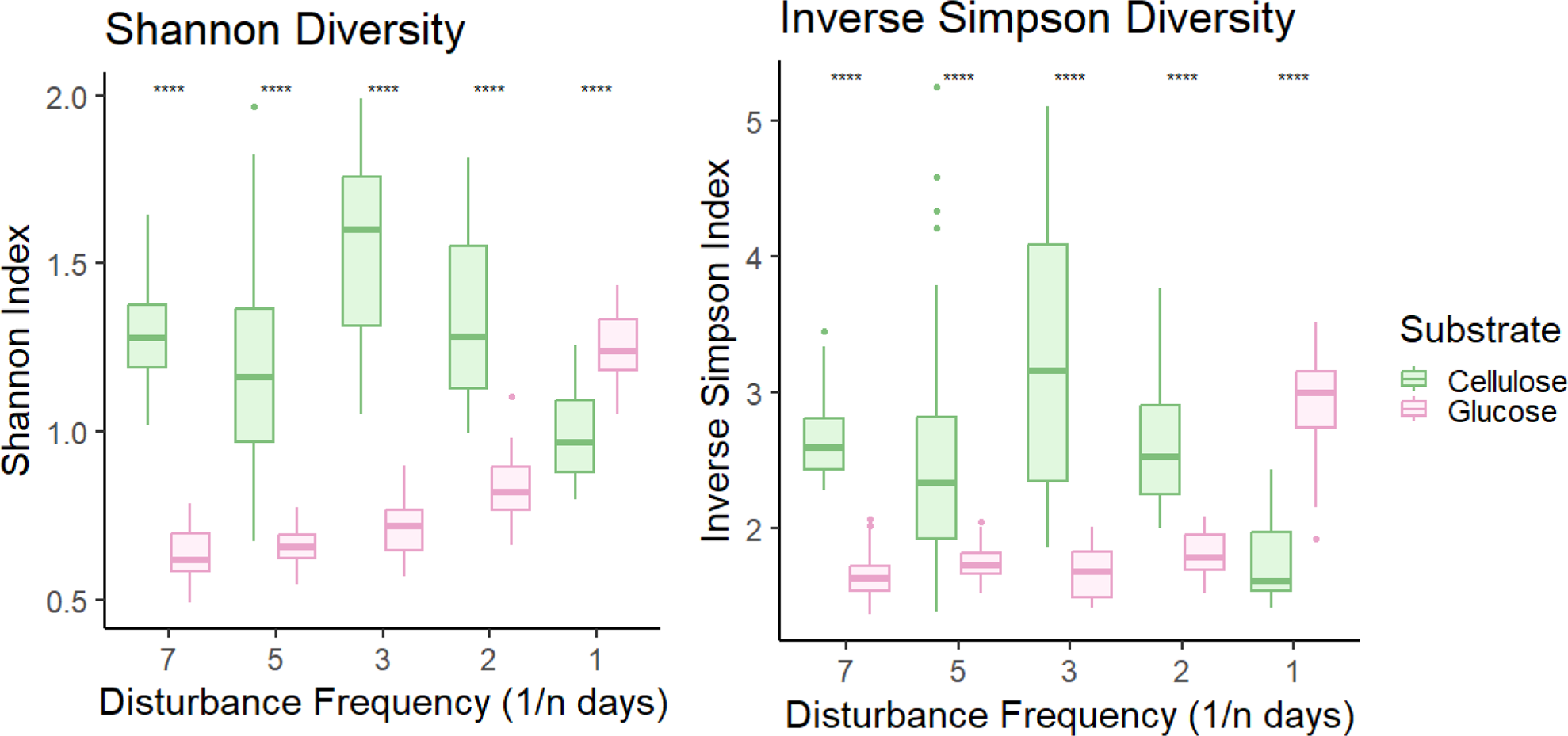
Community diversity varies across substrate and disturbance frequency. Shannon diversity and Inverse Simpson diversity (Y-axis) change across disturbance frequency (X-axis) and vary by substrate.

We also measured Pielou’s evenness and Menhinick’s richness to supplement our alpha-diversity analysis (Supplemental Fig. 3). Samples grown on glucose are more even than cellulose samples at disturbance frequencies 1/7, 1/5, and 1/1 days. In terms of richness, we observed a greater number of unique genera for cellulose samples in nearly all disturbance frequencies, except for 1/2 days where there is a nonsignificant difference.

## Discussion

In this study, we aimed to address how substrate and disturbance frequency interact to shape microbial community structure. We found that substrate is a main driver of the communities’ response to disturbance. We demonstrated that diversity peaks at the intermediate disturbance frequency, 1/3 days, when the community is grown on cellulose, a recalcitrant carbon source. However, community diversity peaks at the highest disturbance frequency, 1/7 days, when the community is grown on glucose, a labile carbon source. The results of this work show how community response to disturbances can be impacted by the substrate they are grown in and contributes to our understanding of how environmental factors interact with disturbances to impact bacterial communities.

### Diversity-Disturbance Relationships

Our community displays a different Diversity-Disturbance Relationship (DDR) depending on the substrate it is grown on. When grown on cellulose, the community displays a unimodal curve (Fig. 2) that fits with predictions of the Intermediate Disturbance Hypothesis, which posits that diversity peaks at intermediate disturbances ^5^. However, the IDH has been found to be an inadequate framework as studies across a variety of ecosystems have found many divergent types of DDRs ^6,31^. Our findings also demonstrate the incompleteness of the IDH; communities grown on glucose display a non-unimodal DDR. Differing DDRs resulting from the same experimental system have been observed before ^18^. Hall et al. 2012 manipulated disturbance intensity (the proportion of cells they moved) and used a simpler one-species community – exploiting the ability of *Pseudomonas fluorescens* to exhibit distinct morphotypes based on access to oxygen. They found a flat, monotonically increasing, or unimodal DDR depending on the disturbance intensity ^18^.

Other experiments have found a variety of DDRs, including a U-shaped DDR ^32^. A model of a two-member community, based on experimental observations, consistently found unimodal DDR, although the exact shape changed with time ^33^. A more recent model of a two-member community displayed multimodality ^34^. One potential reason we did not observe a unimodal DDR with our glucose treatment could be because we did not have a disturbance regime that was frequent enough to result in a population bottleneck. If the disturbances were so frequent that no, or very few, microbes were being passaged each time, then we might expect the diversity of the glucose communities to decrease.

### Disturbance disrupts community composition

Following their assigned disturbance regime treatment, we sampled our experimental communities over the course of a week to evaluate how disturbance frequencies may impact community assembly. In the intermediate disturbance frequency for cellulose treatment (1/3 days), *Cellvibrio* is typically abundant, before being replaced by other taxa. This succession resembles what Lewin et al. 2022 reported. However, at lower frequencies (1/7 and 1/5 days) *Lacunisphaera* was also found to be abundant, and at higher frequencies (1/2 days and 1/1 days) *Asticcacaulis* increases in relative abundance. Notably, *Cellvibrio* starts at lower relative abundance before increasing in the high frequency disturbance treatments (1/2 and 1/3 days).

It is important to note that *Lacunisphaera* was not identified in Lewin’s work. *Lacunisphaera* spp., which belong to the phylum Verrucomicrobia phylum do not have any reported have cellulolytic activity, although one isolate has been described to use a variety of carbon sources ^35^. An *Asticcacaulis* ASV was abundant in our high frequency cellulose treatments and an *Asticcacaulis* OTU was found in Lewin et al. 2016 ^13^. *Asticaccaulis* has been found in other lignocellulolytic communities including communities derived from wood or forest soil^36,37^.

Communities grown in glucose did not display obvious assembly patterns at most disturbance frequencies. We identified two abundant *Pseudomonas* ASVs in the glucose substrate treatments. We cannot determine if these represent different populations, but they appear to have different dynamics across disturbance frequencies. As our study was limited to 16S rRNA gene amplicon sequencing, we cannot determine what mechanisms led to the abundance of *Pseudomonas* ASVs in the glucose samples. *Pseudomonas* is a common environmental microbe, known best as a soil-dweller or member of the rhizosphere microbiome ^38^. As enteric bacteria, the *Pseudomonas* ASVs likely have faster growth rates than other ASVs in these communities. Enrichment for copiotrophs when growth substrate is supplemented with labile carbon has been observed in a previous study ^39^. Additionally, *Pseudomonas* are known for producing a variety of natural products, including molecules that suppress competing microbes ^40,41^. This may be one explanation for how it came to dominate the glucose samples.

The two *Psuedomonas* ASVs dominated community composition in most disturbance frequencies for communities grown in glucose, except for the highest frequency treatment which also had highly abundant *Cohnella* ASV. An isolate from this genus has shown cellulolytic ability ^42,43^. Although we cannot explain why it is abundant in the high frequency glucose samples, the same ASV is also found in the intermediate frequency of our cellulose samples, which matches the report of Lewin et al. 22 that found *Cohnella* to be positively associated with cellulose degradation ^14^.

The different DDRs and successional patterns we observe are likely due to the differing interactions between microbes in the two substrates we considered. Cellulose is a recalcitrant substrate that must be cleaved into cellobiose (a glucose dimer) which is transported into the cell before being cleaved into glucose ^44^. *Cellvibrio* is likely the dominant cellulose-degrader in this microcosm ^13,14^. In order to degrade cellulose, *Cellvibrio* produces extracellular endoglucanases and exoglucanases that liberate cellobiose from the cellulose polymer ^45^. Excess cellobiose molecules are likely what feeds the remaining community. Thus, non-cellulolytic organisms cannot immediately consume carbon in our cellulose treatments and must wait for cellulose-degraders to enrich the media with labile carbon. In contrast, glucose is labile, and thus competition is likely a much stronger driving force in community dynamics. Given that we used 16S rRNA gene amplicon sequencing, we cannot make definitive conclusions about the type of interactions in our community. However, substrate complexity is known to influence bacterial interactions. For example, a synergistic interaction found in co-cocultures of *Citrobacter freundii* and *Sphingobacterium miltivorum* on carboxymethyl-cellulose, xylan, lignin or wheat straw was lost when the pair was grown on glucose ^46^.

## Conclusion

Here, we have demonstrated that communities will respond differently to the same disturbance regime, when grown on substrates of varying complexity. We observe a unimodal DDR when communities are grown on cellulose, a recalcitrant substrate. When grown on glucose, however, we observed a monotonically increasing DDR. Although substrate is a strong predictor for community composition, communities further cluster by disturbance frequency, and successional dynamics differ between disturbance treatments for the same substrate. These results suggest that the range of DDRs we observe across different microbial systems may be due to the nutritional resources microbial communities can access and the interactions between bacteria and their environment.

## Acknowledgements

We are thankful to Soleil E. Young for her comments on drafts of this paper.

This work was funded by the DOE Great Lakes Bioenergy Research Center (DOE BER Office of Science DE-FC02-07ER64494). Funding for D.Q.H. was provided by the National Institutes of Health (National Research Service award T32 GM007215), and SciMed Graduate Research Scholars Fellowship UW-Madison.

---

**Supplemental Table 1.** Media recipe.

|  |  |  |  |  |  |  |  |
| --- | --- | --- | --- | --- | --- | --- | --- |
| M63 Media |  |  | Five-X |  |  | Trace Element Solution SPV-4 |  |
| Five-X | 200 mL |  | Potassium phosphate dibasic K2HPO4 - anhydrous | 53.6 g |  | CaCl2·2H2O | 4 g |
| 1 M MgSO4 | 1 mL |  | Potassium phosphate monobasic KH2PO4 - anhydrous | 26.2 g |  | FeCl2·6H2O | 1.1 g |
| Thiamin (B1) | 1 mL |  | Ammonium sulfate (NH4)2SO4 | 10 g |  | MnSO4·H2O | 0.22 g |
| Trace elements | 5 mL |  | Iron Sulfate (1mg/mL in 0.01 M HCL) | 2.6 mL |  | ZnCl2 | 0.1 g |
| Distilled water | 1 mL |  | Distilled water | 1 L |  | CUSO4·5H2O | 0.04 g |
| Carbon source | 5g |  |  |  |  | CoCl2·6H2O | 0.04 g |
|  |  |  |  |  |  | Na2MoO4·2H2O | 0.025 g |
|  |  |  |  |  |  | Na2B4O7·10H2O | 0.1 g |
|  |  |  |  |  |  | Distilled water | 1 L |

**Supplemental Figure 1.**
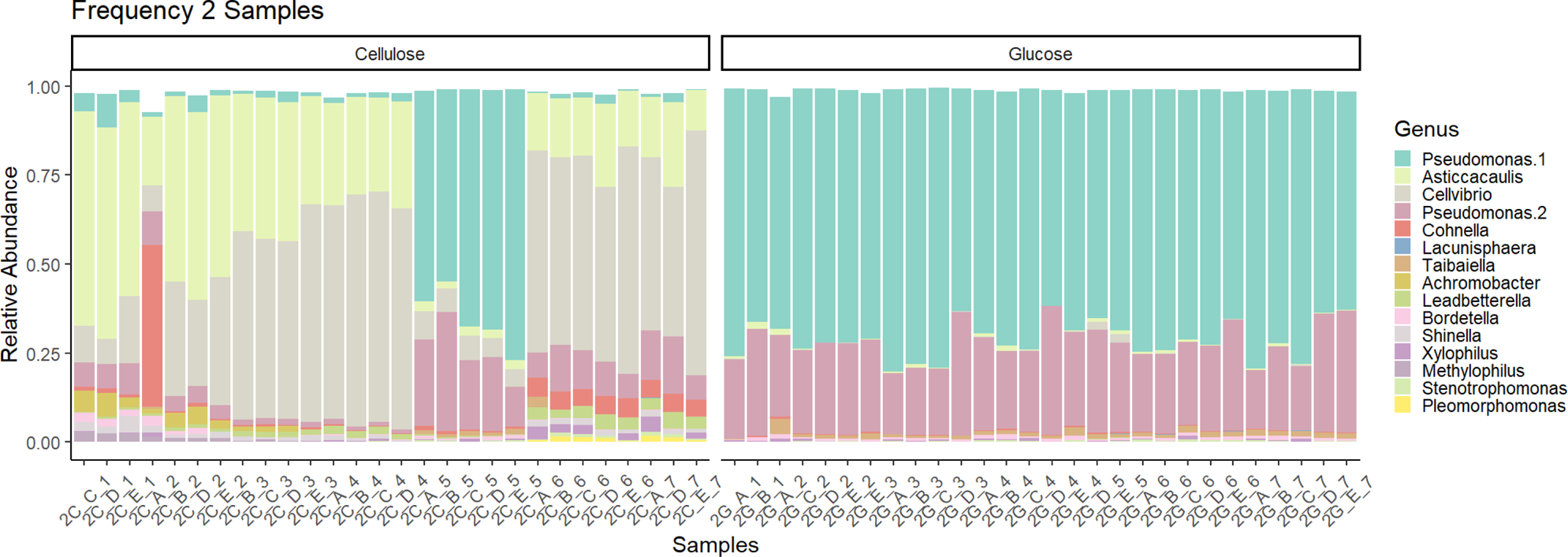
Relative abundance plot of samples from the disturbance frequency 1/2 treatment. Samples from day 5 of the cellulose treatment were removed from analysis, as we suspect they were mislabeled or mis-pipetted at some point prior to sequencing.

**Supplemental Figure 2.**
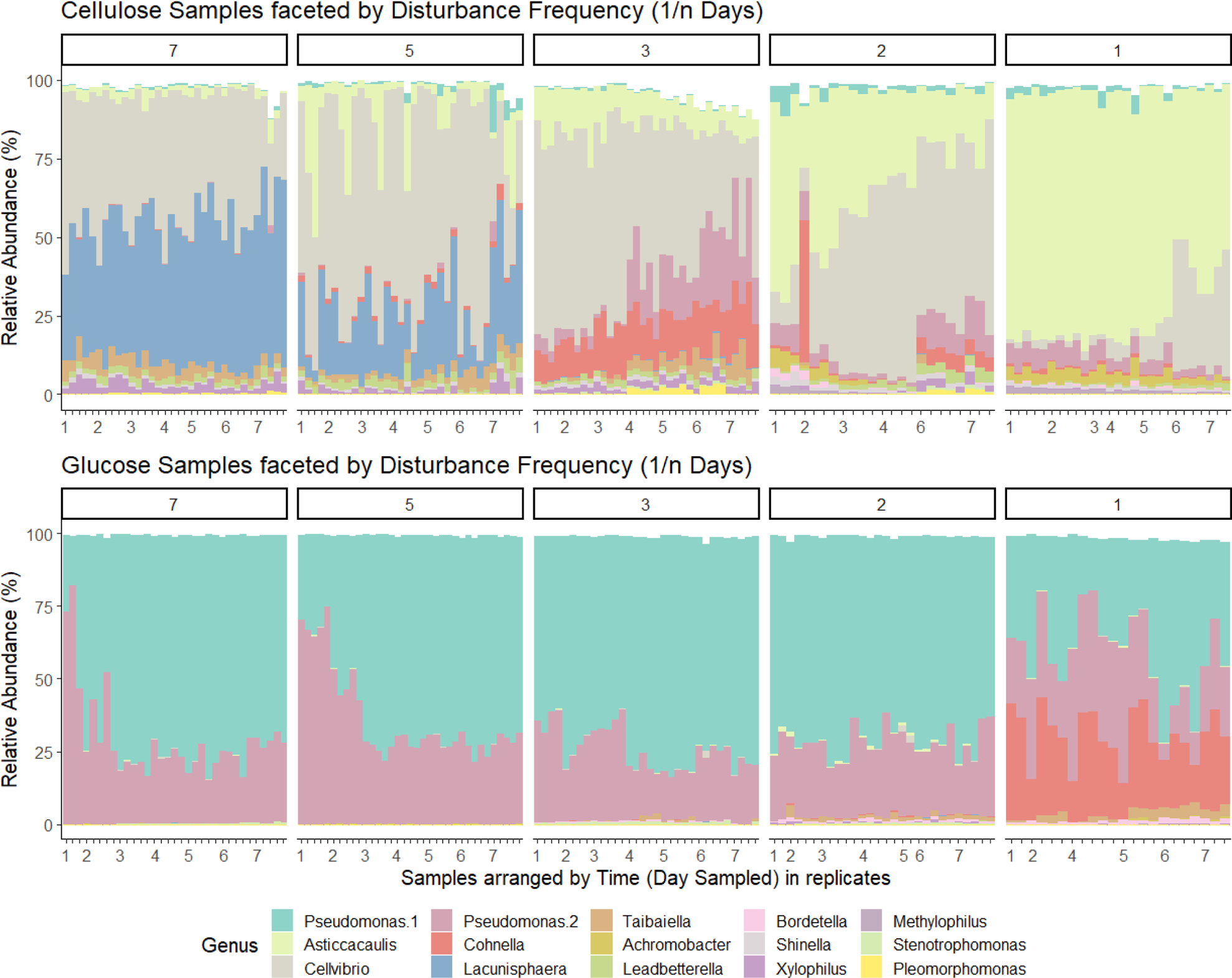
Bar graph of community composition based on relative abundance of 16S rRNA gene amplicon sequencing of the top 15 most abundant ASVs and their assigned genus. Plots are faceted by disturbance frequency (1/n days). Within each facet, samples are grouped by time sampled to observe any compositional dynamics.

**Supplemental Figure 3.**
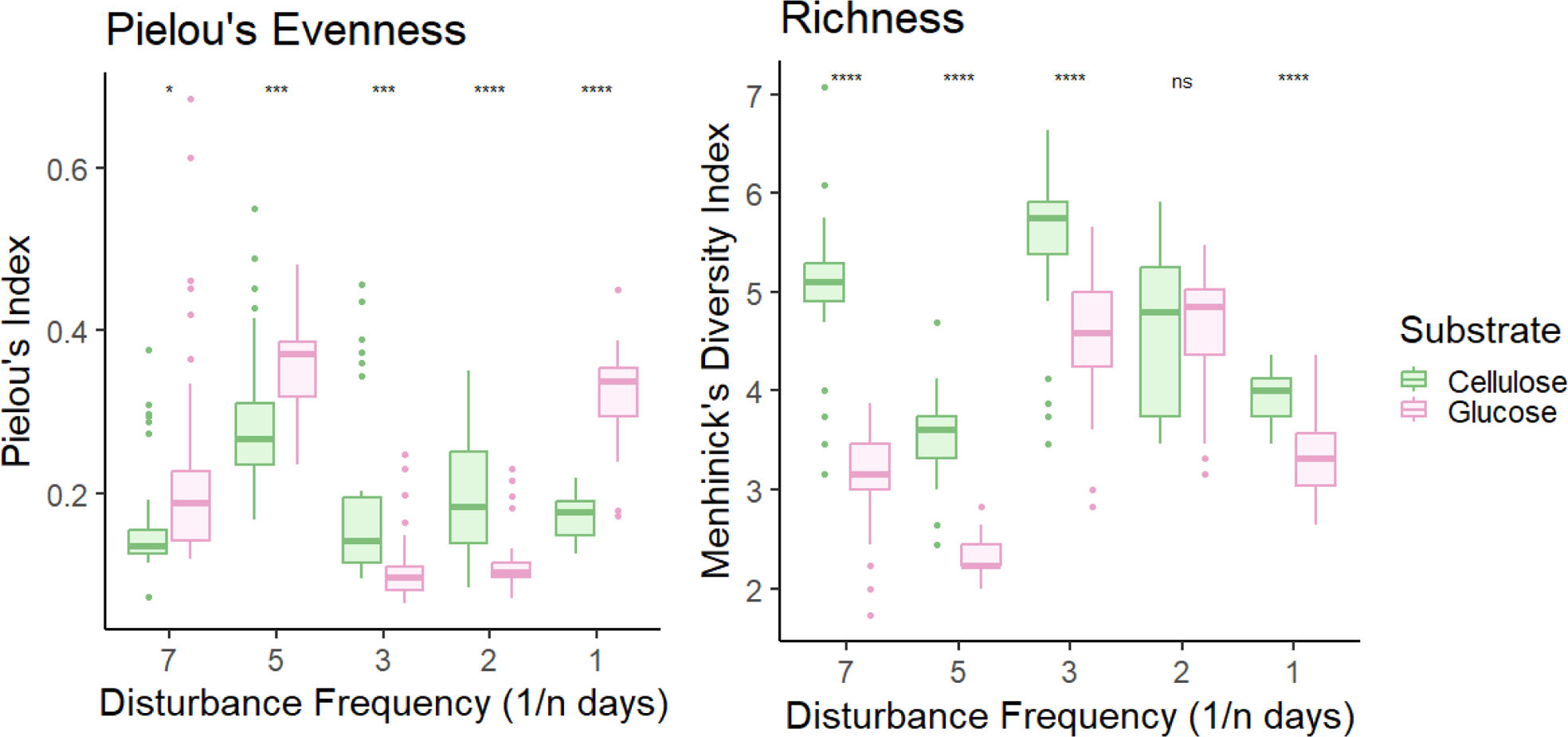
Pielou’s evenness and Menhinick’s richness indices across disturbance frequency. Significance values represent t-test comparing cellulose and glucose samples, grouped by Disturbance Frequency.

**Supplemental Figure 4.**
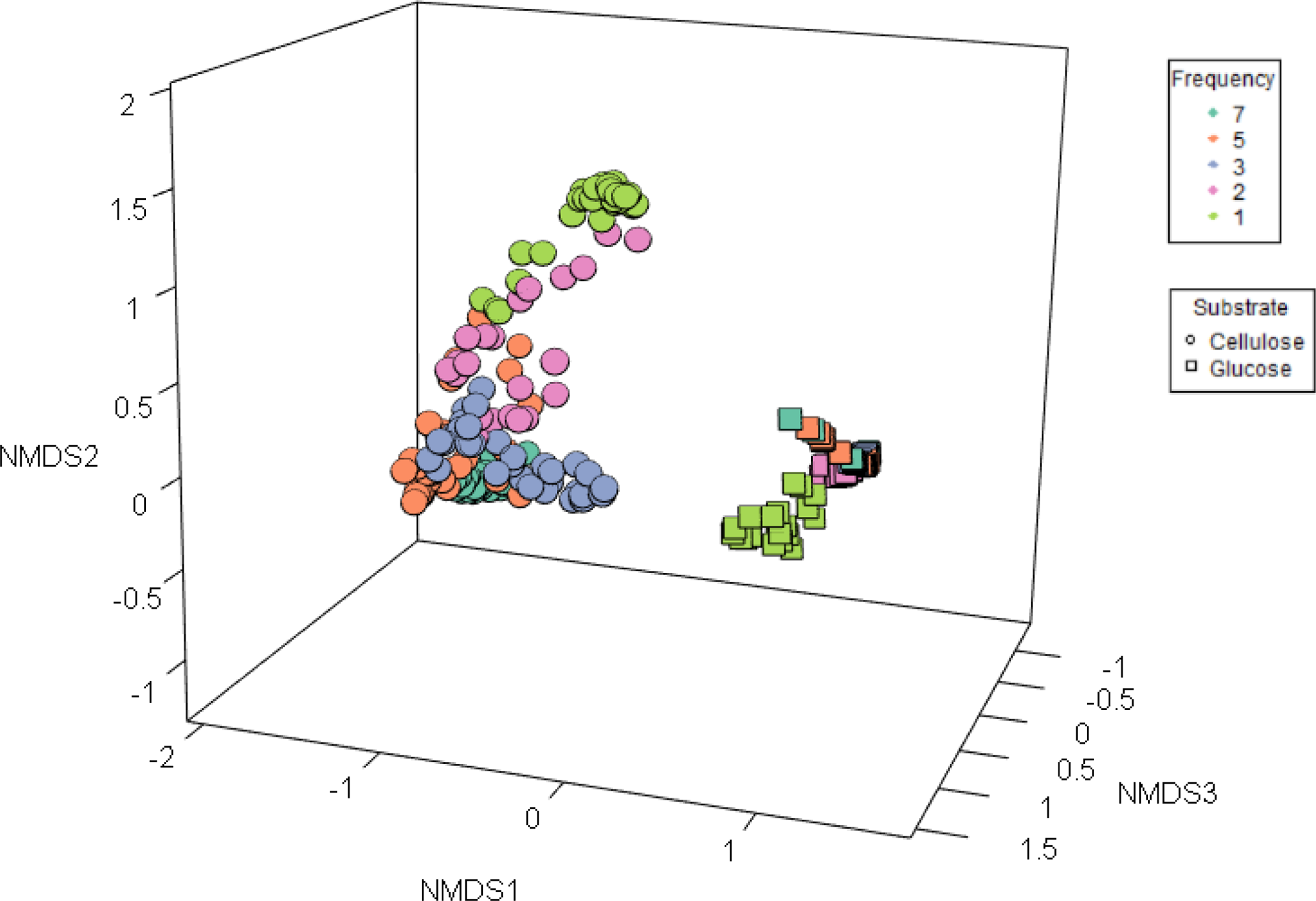
A view of a 3D NMDS plot of community composition. Dissimilarity matrix was calculated using Bray-Curtis dissimilarity. Communities grown in cellulose are shown as circles, and communities grown in glucose are shown as squares. Disturbance frequency is marked by color.

**Supplemental File 1.** NCBI accession numbers and metadata of 16S rRNA gene sequencing data generated for this work.

